# Garlic-Derived Allicin Modulates Expression of the SARS-CoV-2 Spike Receptor-Binding Domain in Human Splenic Fibroblasts

**DOI:** 10.1101/2025.06.10.659002

**Authors:** Ashpreet Kaur, Barnabe D. Assogba

## Abstract

Allicin, a natural sulfur-rich compound, is formed when garlic is crushed or chopped. It has been used for centuries as a natural defence against bacteria and fungi. However, we do not fully understand how well it works against viruses, such as SARS-CoV-2. In this study, we investigated the effect of allicin on the expression of the SARS-CoV-2 receptor-binding domain of the spike protein in human primary splenic fibroblasts. These cells were transfected with a plasmid encoding RBD-S protein fused to a superfolder green fluorescent protein (sfGFP), a bright, stable marker. When cells were treated with 50□µM allicin, either 30 min before or 6 h after transfection, a pronounced drop in fluorescence was observed, indicating reduced spike protein expression. Interestingly, the cell morphology, growth, and behavior remained normal, indicating that allicin could minimize spike protein levels without being toxic to the cells. These results open the door to the use of allicin as a gentle, natural antiviral agent and raise critical questions regarding the regulation of protein expression after transcription. Further research is required to understand these effects better.

## Introduction

Garlic (*Allium sativum*) has been revered since ancient times, not only for its culinary applications as a pungent spice but also for its numerous health benefits (1). As a member of the Allium family of plants, garlic is a treasure trove of bioactive compounds including alliin, ajoene, saponins, and other sulfur compounds (2,3). Allicin (diallyl thiosulfinate) has been identified as the most pharmacologically active compound and extensively studied (4). Allicin is not found naturally in fresh garlic cloves as an active ingredient but is enzymatically generated upon crushing, chopping, or mastication of garlic. This mechanical trauma results in the degradation of alliinase, which rapidly converts the precursor molecule alliin (S-allyl-l- cysteine sulfoxide) into allicin(5–7). This unstable, sulfur-containing substance not only provides a garlicky odor to garlic but also creates most of its therapeutic action (1).

Extensive studies have demonstrated the broad-spectrum biological activities of allicin, including its antimicrobial, antifungal, anti-inflammatory, antioxidant, and anticancer properties (8,9).Allicin has also been studied for its beneficial effects on cardiovascular diseases, specifically through the regulation of blood pressure and lipid concentration (10). Nevertheless, despite its promising traits, the antiviral activity of allicin has been relatively less explored, especially against newly emerging global health concerns, such as the COVID- 19 pandemic(11,12).

SARS-CoV-2, the COVID-19 pathogen, uses its spike (S) protein to infect host cells by binding to the ACE2 receptor, which is expressed extremely highly in respiratory, cardiac, and intestinal tissues (13). Thus, spike proteins have become leading candidates for vaccine and antiviral drug development (14–17). Apart from its role in infection, current studies have also focused on the role of the spike protein in post-COVID neurological symptoms (18) and cardiovascular disease in the absence of active viral replication(19–21). These findings highlight the need for solutions that can downregulate spike proteins or neutralize their downstream effects.

In the present study, we explored whether allicin could downregulate the expression of subunit 1, specifically the receptor-binding domain (RBD) of the SARS-CoV-2 spike protein in human primary splenic fibroblasts. To investigate this, cells were transfected with pcDNA3-SARS-CoV-2-S-RBD-sfGFP, a plasmid that harbors the receptor-binding domain of the spike gene with superfold-GFP (sfGFP) fused, using Lipofectamine 3000 and treated with 50□µM allicin 30 min before (pre-treatment) or 6 h after (post-treatment) transduction. This concentration was chosen based on previous studies that have demonstrated biological activity with no cytotoxicity (22–25). The 6-hour post-transfection time point was selected based on our lab optimization, which showed maximal transfection efficiency at this time point (26).

Quantification of superfolder green fluorescent protein (sfGFP) intensity by fluorescence microscopy revealed that pre-and post-treatment with allicin significantly inhibited RBD-spike protein expression. This was a repeatable effect between replicates, and was not due to changes in cell morphology or viability.

These results suggest a novel role for allicin in modulating the expression of the RBD-spike protein at the cellular level. While more research is needed to determine the precise mechanisms involved and whether the full-length spike protein is also affected by allicin, it is also of interest to investigate whether this downregulation occurs at the translational or transcriptional level. Collectively, our findings provide a promising foundation for examining the antiviral activity of allicin in future therapeutic regimens.

## Materials and Methods

### Cell Culture

Primary human splenic fibroblasts were cultured in T25 flasks using complete fibroblast medium (Cat# M2267, Cell Biologics), supplemented with 25□mL fetal bovine serum (FBS), 5□mL Antibiotic-Antimycotic Solution, 0.5□mL fibroblast growth factor (FGF), and 0.5□mL hydrocortisone, following the manufacturer’s guidelines. cells were maintained at 37°C in a humidified 5% CO□ incubator. Morphology, behavior, and confluency were routinely monitored under a Leica MC120 HD microscope, and images were captured using LAS EZ software. The medium was regularly refreshed. Once 80–90% confluence was reached, the cells were passaged using 3 mL of 0.05% trypsin-EDTA (Cat# 25300054, Invitrogen) for 5–7 min, followed by neutralization with 10 mL of complete medium. For experimental assays, cells were seeded in three 6-well plates (cat# 353046, Life Sciences) designated for allicin treatment and transfection.

### Allicin Preparation

Allicin (Product Code: HY-N0315, MedChemExpress, New Jersey, USA) was supplied as a 100 mg liquid stock. For experimental use, 16.2 μL of pure stock was diluted in sterile 1X PBS to prepare a 10 mM working solution. A 50 µM treatment solution was freshly prepared before each experiment by adding 60 µL of 10 mM stock to 11.94 mL of serum-free medium. All dilutions were performed under sterile conditions. unused stock was wrapped in foil to protect from light and stored at –80°C.

### Allicin Treatment and Transfection

Cells in plates A, B, and C (figure not shown) were cultured for one week to ensure uniform attachment and growth. Plate A: Control (A1–A3) received 2□mL serum-free medium for 30□min at 37□°C, followed by complete medium. Wells A4–A6 were treated with 2 mL of 50 µM allicin, incubated for 30 min, washed with PBS, and then returned to complete medium. Plate B: Wells B1–B3 were transfected with 1.75□mL serum-free medium and 250□µL of transfection mix containing pcDNA3-SARS-CoV-2-S-RBD-sfGFP plasmid and Lipofectamine 3000. Wells B4–B6 received the same allicin pre-treatment as wells A4–A6 before transfection. Plate C: Wells C1–C3 received 50□µM allicin 6□h after mock exposure of the medium. Wells C4–C6 were transfected as in Plate B and treated with allicin 6□h post- transfection. Transfection complexes were prepared in two tubes: Tube 1:18.75□µL plasmid DNA + 6□µL P3000 reagent in 725.25□µL serum-free medium. Tube 2:6□µL Lipofectamine 3000 in 744□µL of serum-free medium. After 10–15 min of incubation, tube 1 was added to tube 2 and gently mixed, and 250 µL was added to each well. All plates were incubated for 24 h prior to analysis, as determined by optimization.

## Results

### Allicin Does Not Affect Morphology or Viability of Splenic Fibroblasts

To evaluate the cytocompatibility of Allicin, we first examined its effects on the morphology of human splenic fibroblast cells (HSFCs). After 24 h of treatment with 50 µM allicin, the cells retained their classical elongated, spindle-like morphology and adhered uniformly in a monolayer, closely resembling that of untreated controls (Figure 1). No signs of cytotoxicity such as cell rounding or detachment were observed. These findings confirmed that the chosen concentration of allicin did not compromise cell viability or structural integrity under the experimental conditions.

**Figure 1.**
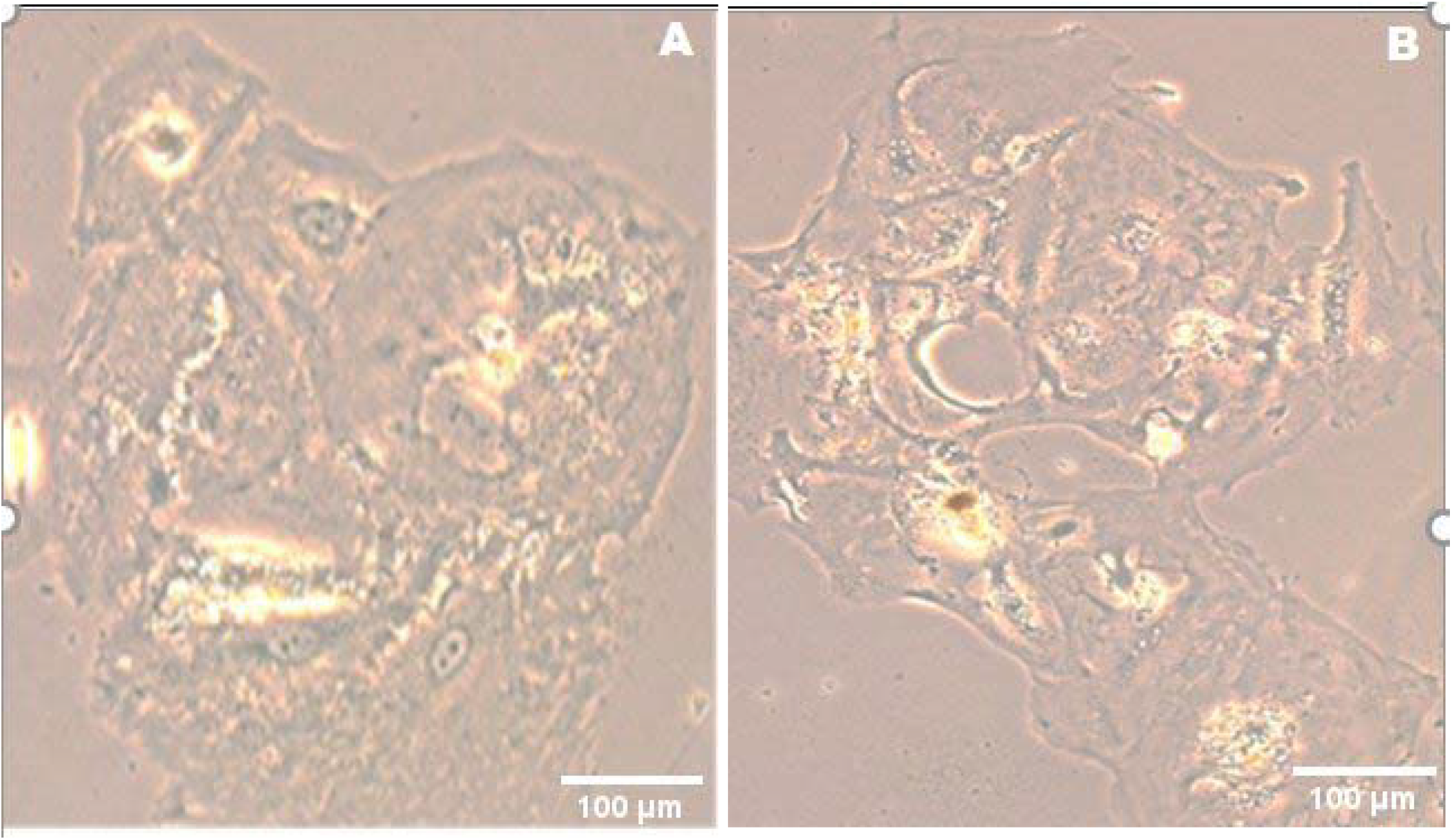
Morphology of human primary splenic fibroblast cells (HSFCs) following 24-hour culture in control and allicin-treated conditions. The micrographs show human primary splenic fibroblast cells cultured in (A) mock (untreated control) and (B) allicin-treated (50 µM) conditions in a humidified atmosphere of 5% CO_2_ at 37 °C. Cells from both groups exhibited classical fibroblast-like morphology, characterized by elongated, spindle-shaped cell bodies and cytoplasmic processes with uniform monolayer growth. There was no observable difference in cell shape, cell density, and adherence between the mock and allicin-treated groups after 24 hours of incubation, indicating that 50□µM allicin does not affect the basal morphology or viability of HSFCs under the studied conditions: scale bar, 100 µm.

### Efficient Spike-RBD-GFP Transfection Observed in HSFCs

To monitor spike protein expression, HSFCs were transfected with the plasmid pcDNA3- SARS-CoV-2-S-RBD-sfGFP using Lipofectamine 3000. Fluorescence microscopy at 24□h post-transfection showed strong sfGFP expression in the plasma membrane and perinuclear regions of transfected cells (Figure 2, Panel 2). In contrast, mock-transfected cells (Figures 1A and 2B) displayed no sfGFP signal, confirming the specificity of transgene expression. Although minor signs of transient lipofection-induced stress (e.g., mild cell rounding) were noted, the overall cell morphology remained preserved. Quantitative fluorescence analysis was performed using four randomly selected transfected cells to assess the subcellular distribution of the signals. Average GFP intensity was significantly higher in the nuclear region (35.78□±□13.83) compared to the plasma membrane (10.21□±□3.95) (Figure 3), suggesting that the RBD-sfGFP fusion protein may preferentially accumulate near or within the nucleus.

**Figure 2.**
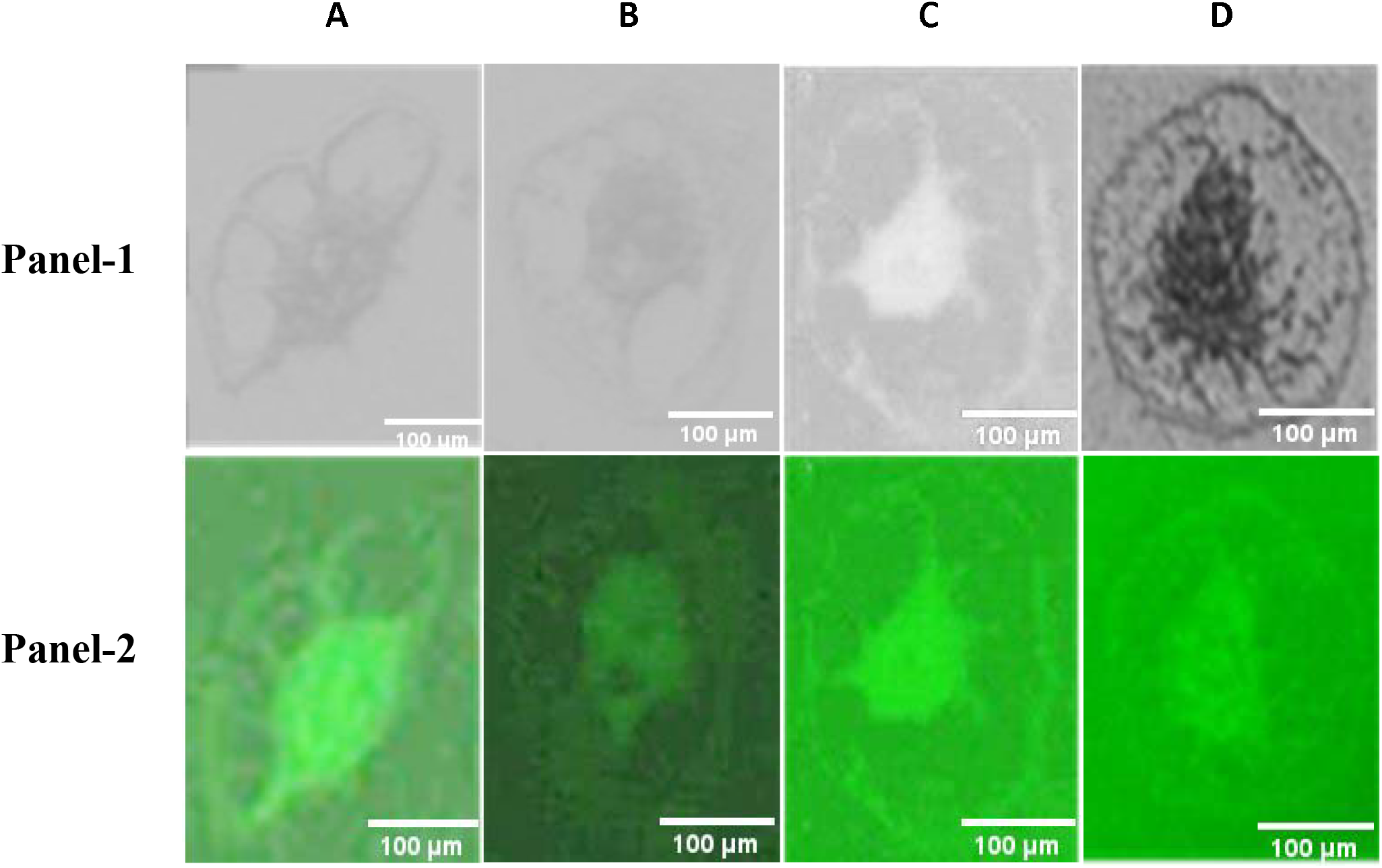
Morphological changes and spike-GFP expression in human primary splenic fibroblasts (HSFCs) following transfection by Lipofectamine 3000. Panel 1 presents the HSFCs under brightfield microscopy after mock transfections, showing the typical fibroblast morphology, with cells elongated and spindle-shaped, growing on an adherent monolayer. The corresponding GFP fluorescence image (A2) shows that there was no background signal. Twenty-four hours post-transfection, Panel 2 shows the HSFCs after transfection with spike-GFP plasmid via the Lipofectamine 3000. Distinctive morphological changes happened in these cells, including rounding and detachment, indicating stress from the transfection. Panel B2 shows the same field under a fluorescence microscope, exhibiting strong green fluorescence in the cytoplasm and perinuclear location, which confirms the strong expression of the spike-GFP fusion protein. These images bear testimony to the consequences of lipofection on primary HSFC morphology and the successful application of the transfection protocol.

**Figure 3.**
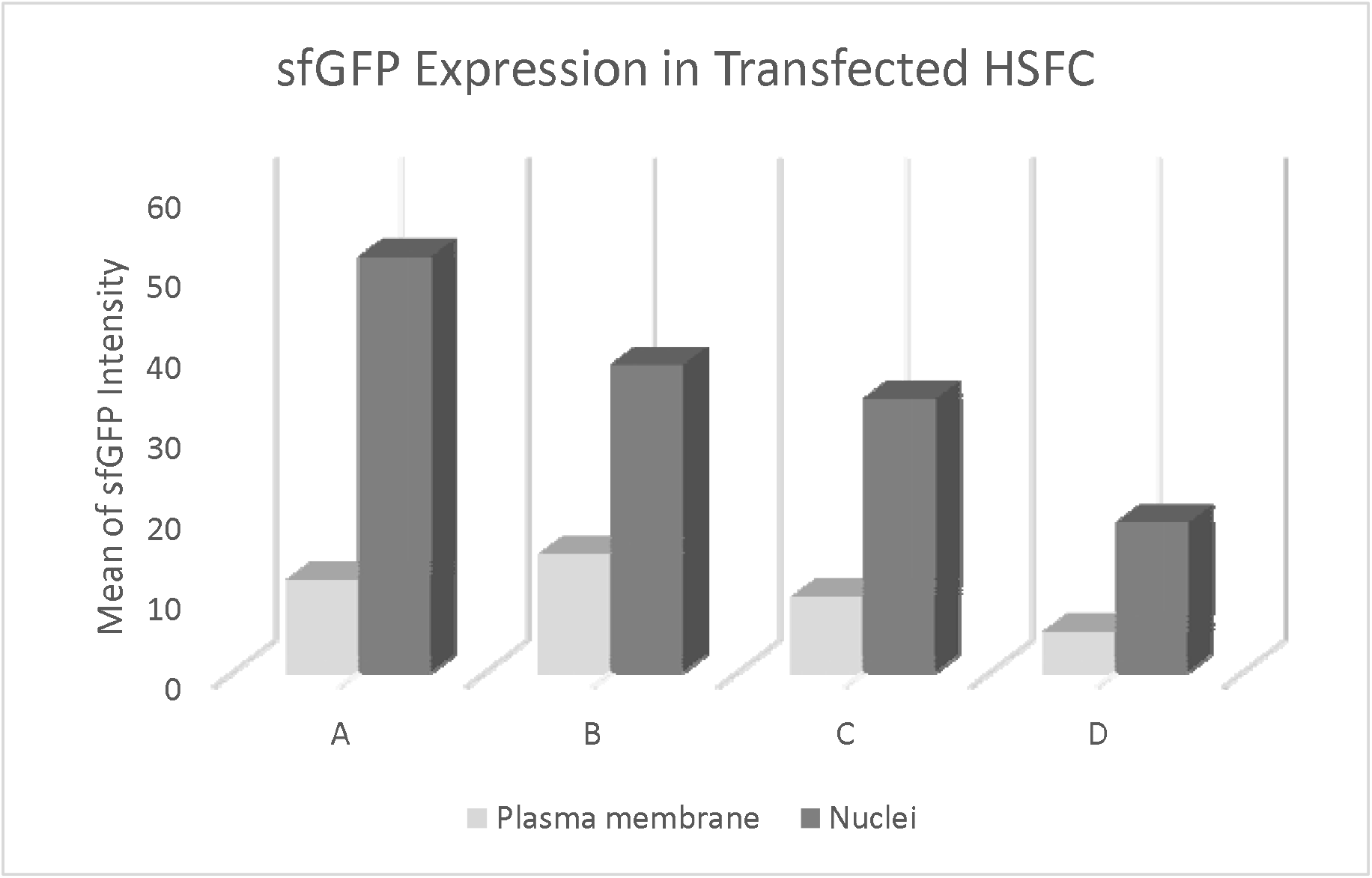
Quantification of sfGFP fluorescence intensity at the plasma membrane and nuclei of transfected HSFCs using ImageJ. The fluorescence intensity of the SARS-CoV-2 spike protein-sfGFP signal was measured using ImageJ in four randomly selected transfected cells (A–D). The mean sfGFP intensity at the plasma membrane was 10.21 ± 3.95, while the mean intensity within the nuclei was significantly higher at 35.78 ± 13.83. These results indicate predominant nuclear accumulation of the sfGFP-tagged protein despite the membrane localization of the spike fusion construct.

### Allicin Strongly Suppresses Spike Protein Expression Regardless of Timing

To test whether allicin could modulate spike protein expression, we compared GFP expression across different treatment groups. Fluorescence microscopy revealed that cells transfected without allicin (B1–B3) showed strong GFP expression, with a transfection efficiency of 96% (Figure 4). However, cells treated with allicin either 30 min before (B4– B6) or 6 h after (C4–C6) transfection displayed a dramatic reduction in the GFP signal, with expression dropping to as low as 4–7% under both pre-and post-treatment conditions. Importantly, in all groups, including those treated with allicin, the morphology of the HSFCs remained consistent, confirming that the downregulation of the GFP signal was not due to cytotoxicity.

**Figure 4.**
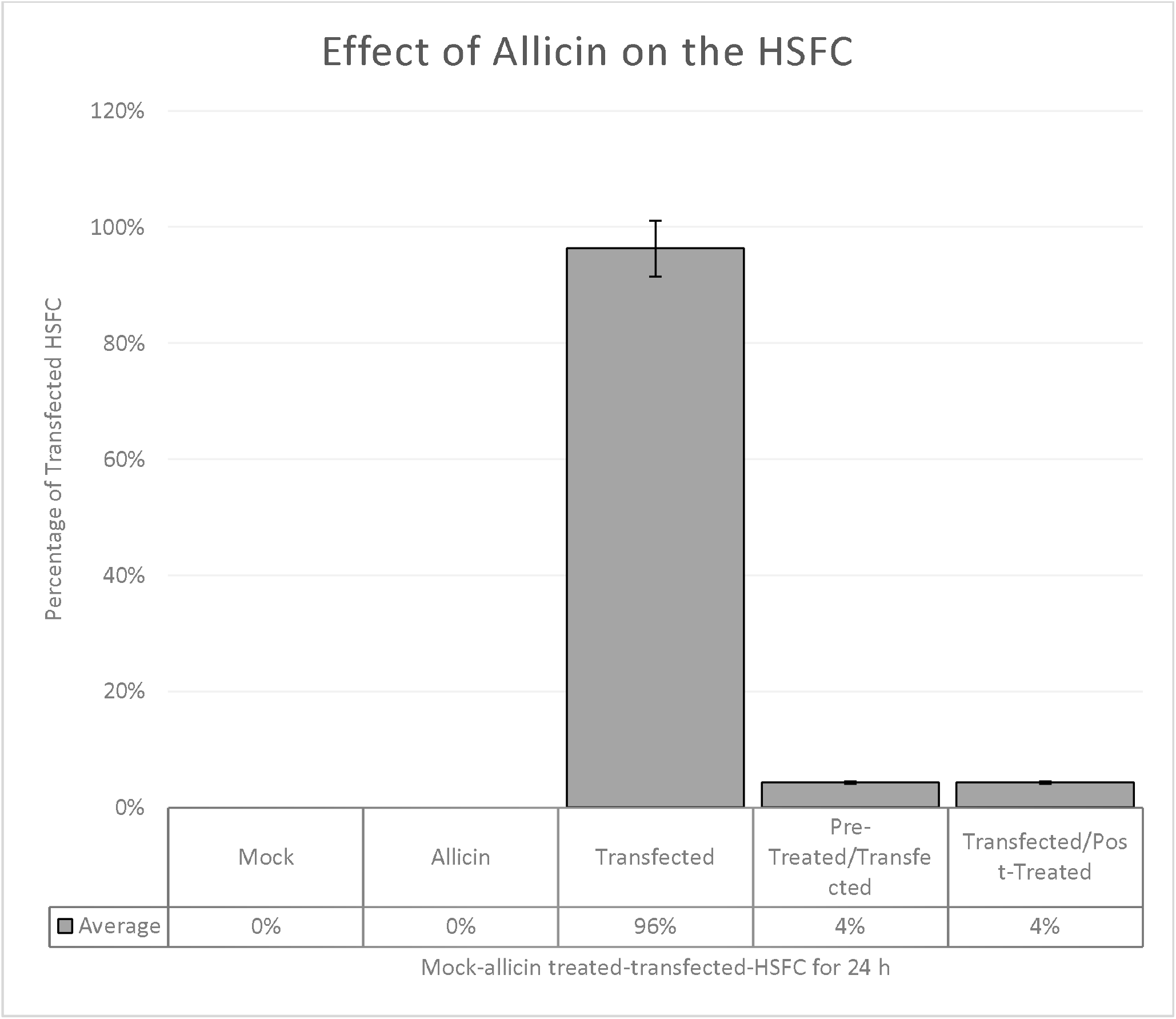
Influence of Allicin pre- and post-treatments on the transfection efficiency of human primary splenic fibroblast cultures. Cells were plated onto three 6-well plates and grouped as follows: Plate A; control (wells A1-A3) and Allicin-treated (50 µM, wells A4-A6); Plate B; transfected (wells B1-B3) and Allicin pre-treated for 30 min, followed by transfection (wells B4-B6); Plate C; transfected (wells C1-C3) and post-treated with Allicin for 6 h after transfection (wells C4-C6). At 24 h post-culture: (1) No morphological differences were observed between control and allicin-only treated groups; (2) High transfection efficiency (96%) was achieved in allicin non-treated cells; (3) No significant differences in transfection efficiency or cellular morphology were noted between pre- and post-allicin treatment groups. These findings suggest that 50 µM allicin does not impair transfection efficiency or alter cellular morphology under the tested conditions. A one-way ANOVA followed by the Tukey post hoc test revealed a significant difference between the groups. The untreated group had a significantly higher transfection efficiency (96.3□±□1.5%) than the Allicin pre- (4.3□±□2.5%) and post- (4.3□±□4.7%) groups (p□<□0.001). There was no difference between the pre-and post- treatment periods. The allicin-only and mock groups were ineffective. When applied either before or after, Allicin at 50 µM significantly lowers transfection.

These results indicated that allicin significantly reduced the expression of the SARS-CoV-2 spike RBD protein in HSFCs. This suppression occurs regardless of whether allicin is administered before or after transfection, suggesting that its mechanism may involve post- transcriptional or translational interference rather than direct effects on transfection efficiency.

## Discussion and Conclusion

This study provides novel insights into the antiviral potential of allicin, a well-known sulfur- containing compound derived from garlic, by demonstrating its ability to suppress the expression of SARS-CoV-2 spike receptor-binding domain (RBD) in human splenic fibroblast cells (HSFCs). This is the first report to explore the effect of allicin on spike protein expression in primary splenic fibroblasts, a cell type relevant to immune system regulation.

Our findings showed that 50□µM allicin markedly reduced spike-RBD-sfGFP fluorescence intensity when administered either before or after transfection, suggesting that its modulatory effects are not time dependent. This reproducible suppression was not associated with any observable cytotoxicity or morphological changes, indicating a selective action on spike protein expression pathways. These observations are in agreement with previous reports that established the ability of allicin to inhibit viral replication and protein expression in other viral systems, including influenza and rhinoviruses, albeit through largely undefined mechanisms.

The preserved cellular morphology across treatment conditions supports the non-cytotoxic nature of the treatment, consistent with previous studies showing that 50 µM allicin is generally well tolerated in human cell lines. This is particularly important given that many antiviral agents exhibit dose-dependent cytotoxicity, thereby limiting their therapeutic potential.

One of the most intriguing outcomes was the similar degree of spike protein suppression observed in both pre-and post-transfection groups. This may imply that the mechanism of action of allicin lies downstream of transcription, possibly affecting mRNA stability, translation initiation, or protein degradation pathways such as the ubiquitin-proteasome system. Allicin interacts with thiol groups on cysteine residues, potentially altering protein folding, stability, or function. Such interactions can destabilize spike protein translation or trigger accelerated degradation. Further mechanistic studies, including transcriptomic and proteomic analyses, are necessary to determine whether allicin interferes at the post- transcriptional, translational, or posttranslational levels.

The subcellular distribution of the spike-RBD-sfGFP fusion protein, which was more concentrated near the nucleus than at the plasma membrane, is in agreement with previous reports that the spike protein localizes to the endoplasmic reticulum-Golgi intermediate compartment (ERGIC) during biosynthesis. Whether Allicin affects intracellular trafficking or folding of the spike protein remains an area of interest.

It is essential to note that although our study employed a plasmid-driven overexpression system and did not involve live virus infection, the ability of allicin to suppress spike protein expression in this controlled context provides foundational proof of concept. This suggests that allicin and its analogs may hold therapeutic promise, either as direct antivirals or adjuncts to existing therapies, by lowering spike-mediated pathogenesis. Given emerging evidence that spike proteins may contribute to post-acute sequelae of SARS-CoV-2 infection (PASC) or “long COVID,” even in the absence of viral replication, the ability to downregulate spike protein levels is clinically relevant.

Collectively, our results provide preliminary but compelling evidence that allicin can selectively and efficiently downregulate SARS-CoV-2 spike RBD protein expression in human splenic fibroblasts, without compromising cell viability. These findings support further investigations of allicin as a natural low-toxicity candidate for complementary antiviral strategies. Future studies should focus on deciphering the underlying molecular mechanisms, extending the findings to full-length spike constructs, and testing more physiologically relevant models, including infection with pseudo-typed or live SARS-CoV-2 under Biosafety Level 3 (BSL-3) conditions.

## Author contributions

Ashpreet Kaur: Conceptualization, Grant preparation, experimental work, data collection, and writing of the original draft.

Barnabe D. Assogba: Project supervision, experimental guidance, Data Analysis, Writing, review, and editing.

## Acknowledgement

This research was generously supported by funding from Kwantlen Polytechnic University (KPU). We express our sincere gratitude to Dr. Barnabe Assogba for providing me with this meaningful project and for their continuous guidance and encouragement throughout the study. We also thank the Office of Research (ORS) for approving the grant and for extending special appreciation to Cathy Parlee for coordinating the procurement of essential lab supplies. We are grateful to laboratory technicians David and Christina for their assistance in buying laboratory supplies. We also thank our laboratory mates for their encouragement during this project. Finally, we acknowledge the Department of Biology and Faculty of Science and Horticulture at KPU for fostering research opportunities and creating an environment that supports undergraduate scientific inquiry.

## Data Access Statement

The data supporting the findings of this study are available from the corresponding author upon reasonable request. Plasmid sequence confirmation data and representative microscopy images were archived at Dr. Assogba’s Cellular and Molecular Biology Laboratory at Kwantlen Polytechnic University.

## Ethics Statement

This study was conducted in accordance with the ethical standards of Kwantlen Polytechnic University (KPU) and the principles outlined in the Declaration of Helsinki. Primary human splenic fibroblasts were obtained from a certified commercial source (Cell Biologics, Chicago, IL, USA) and adhered to all relevant ethical and regulatory guidelines for procurement of human biological material. No direct interaction with human subjects occurred and no identifiable personal information was used. As such, institutional ethics approval was not required for this study. All experimental procedures involving cell culture and handling of genetic material complied with applicable biosafety regulations.

